# Structural basis for sequence context-independent single-stranded DNA cytosine deamination by the bacterial toxin SsdA

**DOI:** 10.1101/2024.09.08.611884

**Authors:** Lulu Yin, Ke Shi, Yanjun Chen, Reuben S. Harris, Hideki Aihara

## Abstract

DNA deaminase toxins are involved in interbacterial antagonism and the generation of genetic diversity in surviving bacterial populations. These enzymes have also been adopted as genome engineering tools. The single-stranded (ss)DNA deaminase SsdA represents the bacterial deaminase toxin family-2 (BaDTF2) and it deaminates ssDNA cytosines with little sequence context dependence, which contrasts with the AID/APOBEC family of sequence-selective ssDNA cytosine deaminases.

Here we report the crystal structure of SsdA in complex with a ssDNA substrate. The structure reveals a unique mode of substrate binding, in which a cluster of aromatic residues of SsdA engages ssDNA in a V-shaped conformation sharply bent across the target cytosine. The bases 5’ or 3’ to the target cytosine are stacked linearly and make few sequence-specific protein contacts, thus explaining the broad substrate selectivity of SsdA. Unexpectedly, SsdA contains a β-amino acid isoaspartate, which is important for enzymatic activity and may contribute to the stability of SsdA as a toxin. Structure-function studies helped to design SsdA mutants active in human cells, which could lead to future applications in genome engineering.

## Introduction

A diverse family of toxins produced by Gram-negative bacteria is delivered to neighboring bacterial cells via the type VI secretion system (T6SS) to inhibit the growth of competing species and provide a growth advantage to the host strain. Such T6SS-delivered toxins include peptidoglycan hydrolases and phospholipases that promote lysis of target cells, membrane disrupting/pore-forming proteins, metallopeptidases, NAD-glycohydrolases, (p)ppApp synthetases, and DNA endonucleases^1^. A recently discovered addition to this superfamily is the bacterial deaminase toxin family (BaDTF) proteins, which catalyze the deamination of cytosine to uracil and cause mutations in the genomic DNA of recipient cells^2,3^. In addition to causing lethal levels of mutation, the activity of BaDTFs may facilitate the evolution of surviving target bacterial populations and contribute to the development of antibiotic resistance^2^.

BaDTF proteins are phylogenetically classified into at least three sub-families. The first class (BaDTF1) is represented by a double-stranded (ds)DNA deaminase from *Burkholderia cenocepacia*, DddA. The deaminase toxin domain of DddA and its orthologs or their evolved variants have been adopted in the CRISPR-free, DddA-derived cytosine base editors (DdCBEs) that enable base editing of mitochondrial, chloroplast, and nuclear DNA targets^3–6^. Recent structural studies have revealed the mode of target engagement by DddA featuring a unique ‘tandem-displacement’ mechanism of base-flipping^7^. The second class (BaDTF2) is represented by a deaminase toxin from *Pseudomonas syringae* designated the single-stranded (ss)DNA deaminase toxin, SsdA^2^ (**Supplementary Figs. S1** and **S2**). In contrast to DddA, which strongly prefers the 5’-TC motif in deaminating double-stranded (ds)DNA substrates, SsdA was reported to preferentially deaminate cytosines in ssDNA with little sequence-context preference^2^. Interestingly, BaDTF2 is evolutionarily more closely related to DYW motif-containing eukaryotic RNA deaminases than to other BaDTFs^8,9^, although SsdA and other BaDTF2 deaminases themselves do not have the C-terminal DYW motif (**Supplementary Figs. S1** and **S3**). The third class (BaDTF3) has been shown to target both ssDNA and dsDNA^2,10^.

The ssDNA-selective cytosine deaminase activity of SsdA is analogous to that of human APOBEC family deaminases, which provide innate antiviral immunity, drive the evolution of human cancers, and are used widely in base editing technologies^11–15^. However, in contrast to the human APOBEC deaminases with TC or CC-restricted editing preferences, SsdA has a much more relaxed target preference by deaminating cytosine preceded by any of the four (T, C, A, or G) bases and even exhibiting residual activity on dsDNA^2^. The structural basis of the broader substrate selectivity for SsdA is unknown. Here we report the crystal structure of SsdA in complex with ssDNA and corroborating biochemical data, which reveal a unique mechanism of substrate DNA recognition.

High-resolution crystallographic data also show that SsdA contains a β-amino acid isoaspartate (isoAsp), which is important for enzymatic activity and may contribute to the stability of the toxin. Based on this structural information, we generated mutant derivatives of SsdA that are active in human cells and may be useful for genome engineering applications.

## Results

### Overall structure of SsdA-ssDNA complex

To understand how SsdA interacts with target DNA, we sought to crystallize the C-terminal toxin domain (Lys259 to Glu409; the residue numbering according to PDB entry 7jtu)^2^ of *P. syringae* SsdA in complex with its preferred DNA substrate. As reported previously^2^, we observed that wildtype SsdA deaminates cytosines in ssDNA with any base at the 5’ (–1) position, although it exhibited a weak context-dependence (**Supplementary Fig. S4**). The deaminase activity was weaker on dsDNA, with a modest preference for purines at the –1 position. Consistent with these results, we obtained crystals of SsdA with a 15-nucleotide (nt) ssDNA substrate containing a 5’-T**C**G target sequence in the middle (**Fig. 1**). SsdA with substitution of the catalytically essential glutamic acid residue (E348A) was used to capture the enzyme-substrate complex. The structure of the SsdA-ssDNA complex was determined by molecular replacement phasing and refined to 1.94-Å resolution with excellent model quality and fit to experimental data (Rwork/Rfree = 16.3/19.0 %, **Table 1**; **Supplementary Fig. S5**). Two SsdA-ssDNA complexes are in the asymmetric unit with essentially the same conformations. The ssDNA bound to SsdA adopts a V-shaped conformation sharply bent away from the protein, with the target 2’- deoxycytidine nucleotide rotated out at the apex of ‘V’ and engaged in the Zn-bound active site. SsdA contains the core structural elements shared among the deaminase superfamily consisting of two α- helices and a layer of β-sheet^8^. The catalytically essential Zn ion is coordinated by a conserved triad motif comprising His346, Cys367, and Cys370 and positioned between the N-terminal ends of the two α-helices. His346 is stacked against the cytosine base in the active site pocket that is lined on the other side by Thr290, a residue conserved among DNA cytosine deaminases. Our SsdA structure in complex with ssDNA shows minimal structural changes from that in complex with the immunity factor/antitoxin (SsdAI) with the root mean square deviation (r.m.s.d.) of 1.28 Å and 0.65 Å for all and the main chain atoms, respectively, although the active site zinc ion was lost in SsdA bound to SsdAI (PDB 7jtu)^2^.

**Figure 1.**
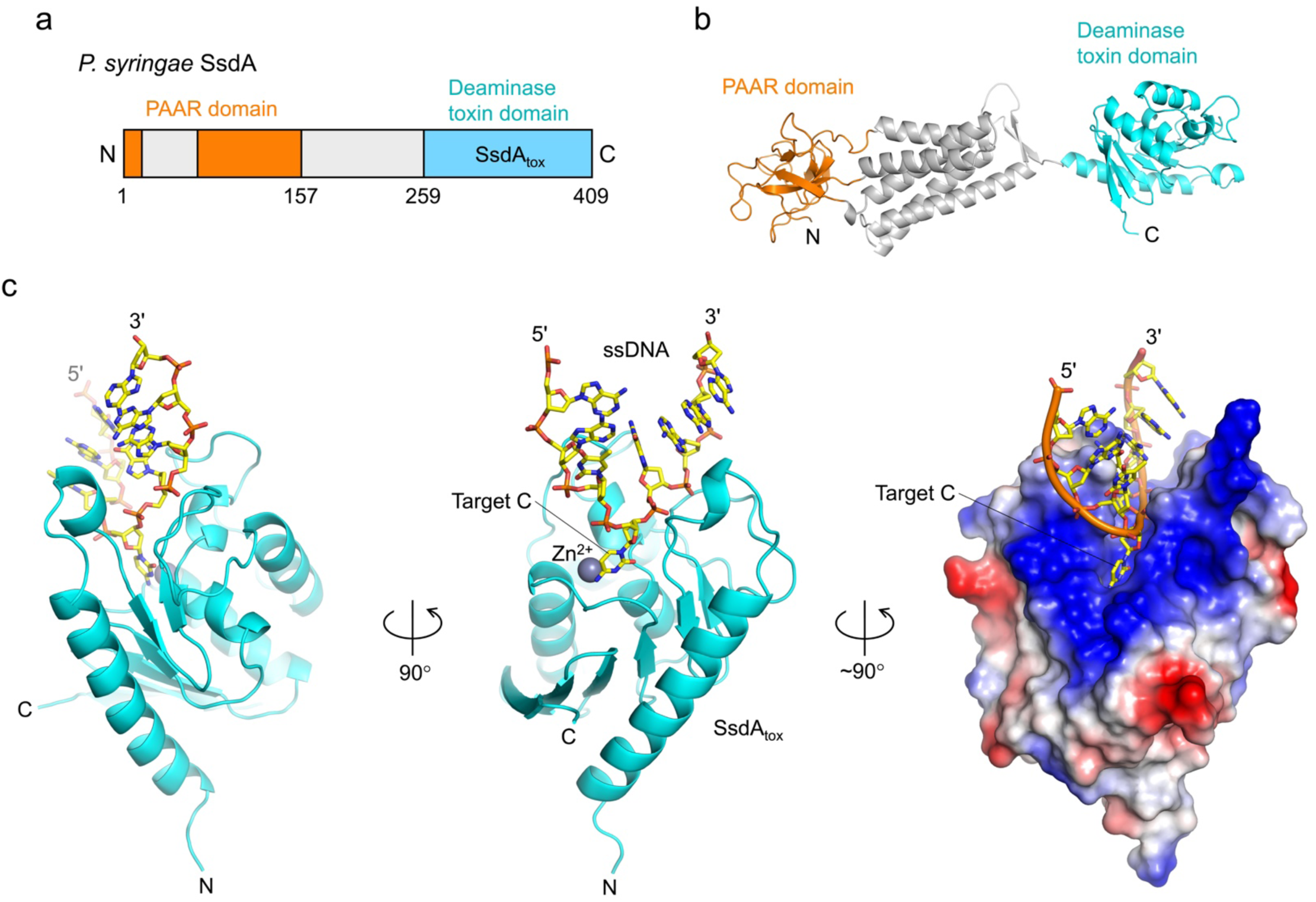
Overall structure of the ssDNA deaminase toxin, SsdA. **a**, A schematic showing the domain organization of *Pseudomonas syringae* SsdA. PAAR: proline-alanine-alanine-arginine. The residue numbering is according to PDB entry 7jtu^2^. **b**, Structure prediction by AlphaFold2^41^ of full-length SsdA. The domains are color-coded as in **a**. **c**, The crystal structure of SsdAtox-ssDNA complex, viewed from three orthogonal orientations. The panel on the right shows the molecular surface of the protein colored according to electrostatic potential (blue: positive, red: negative).

**Table 1.** Crystallographic data collection and refinement statistics.

|  | SsdA | SsdA/ssDNA |
| --- | --- | --- |
| PDB ID | 9C63 | 9C64 |
| **Data collection** |  |  |
| Space group | $P2_12_12_1$ | $P6_1$ |
| Unit cell dimensions |  |  |
| $a, b, c$ (Å) | 52.69, 73.48, 78.06 | 87.92, 87.92, 134.48 |
| Resolution (Å) | 30.14 – 1.33 (1.41 - 1.33) | 66.3 – 1.94 (2.03 - 1.94) |
| $R_{\text{sym}}$ or $R_{\text{merge}}$ | 0.051 (0.796) | 0.152 (1.70) |
| $I / \sigma I$ | 12.8 (1.3) | 7.7 (1.5) |
| Completeness (%) | 93.8 (58.9)* | 94.7 (56.9)** |
| Redundancy | 5.0 (4.0) | 7.1 (7.3) |
| $CC_{1/2}$ | 0.999 (0.602) | 0.997 (0.492) |
| **Refinement** |  |  |
| Resolution (Å) | 30.1 - 1.34 (1.39 - 1.34) | 66.3 - 1.94 (2.01 - 1.94) |
| No. Reflections | 59512 (1376) | 38975 (1226) |
| $R_{\text{work}} / R_{\text{free}}$ | 0.169/0.195 | 0.161/0.190 |
| Non-H atoms | 2793 | 3040 |
| Protein | 2417 | 2399 |
| DNA | N/A | 374 |
| Ligand/ion | 34 | 26 |
| Water | 342 | 241 |
| $B$-factor | 28.97 | 43.70 |
| Protein | 28.02 | 35.91 |
| DNA | N/A | 95.88 |
| Ligand/ion | 24.77 | 33.33 |
| Water | 36.13 | 41.36 |
| Ramachandran plot |  |  |
| Favored (%) | 99.67 | 98.66 |
| Allowed (%) | 0.33 | 1.34 |
| Outliers (%) | 0.00 | 0.00 |
| R.m.s. deviations |  |  |
| Bond lengths (Å) | 0.007 | 0.013 |
| Bond angles (°) | 0.93 | 1.11 |

Statistics for the highest-resolution shell are shown in parentheses.
\* $a^*$ 1.46 Å, $b^*$ 1.34 Å, $c^*$ 1.33 Å
\*\* $0.894a^* - 0.447b^*$ 1.94 Å, $b^*$ 1.94 Å, $c^*$ 2.06 Å

### Mechanism of ssDNA engagement

SsdA engages the sharply bent ssDNA in a deep positively charged groove between two protruding loops. The loop on one side is centered on a short helical segment between Lys284 and Lys287, whereas that on the other side spans between Lys330 and Lys339. The narrow ssDNA-binding groove is most constricted between Val288 and Tyr334, where these residues project into the aperture over the deep active site pocket. The DNA bases on either the 5’ or 3’ side of the target C are each stacked along a linear axis, which are perpendicular to one another (**Fig. 2**). A cluster of aromatic residues of SsdA: His332, Tyr334, Tyr342, and Phe343 serves as a scaffold to shape the ssDNA. Tyr334, in particular, plays a central role in stabilizing the V-shaped DNA conformation – It is π-stacked against the face of the +1 G base to set the direction of the strand downstream (3’) of the target C, while being hydrogen-bonded to –2 A and making a T-shaped aromatic stacking interaction with –3 A to position the upstream (5’) bases. His332 is hydrogen-bonded to the backbone phosphate group of –1 T, also stabilizing the DNA strand upstream of the target C. The thymine base immediately 5’ to the target C (–1 T) is exposed to the solvent and makes no protein contact, which explains the lack of strong sequence context preference of SsdA in deaminating ssDNA substrates.

**Figure 2.**
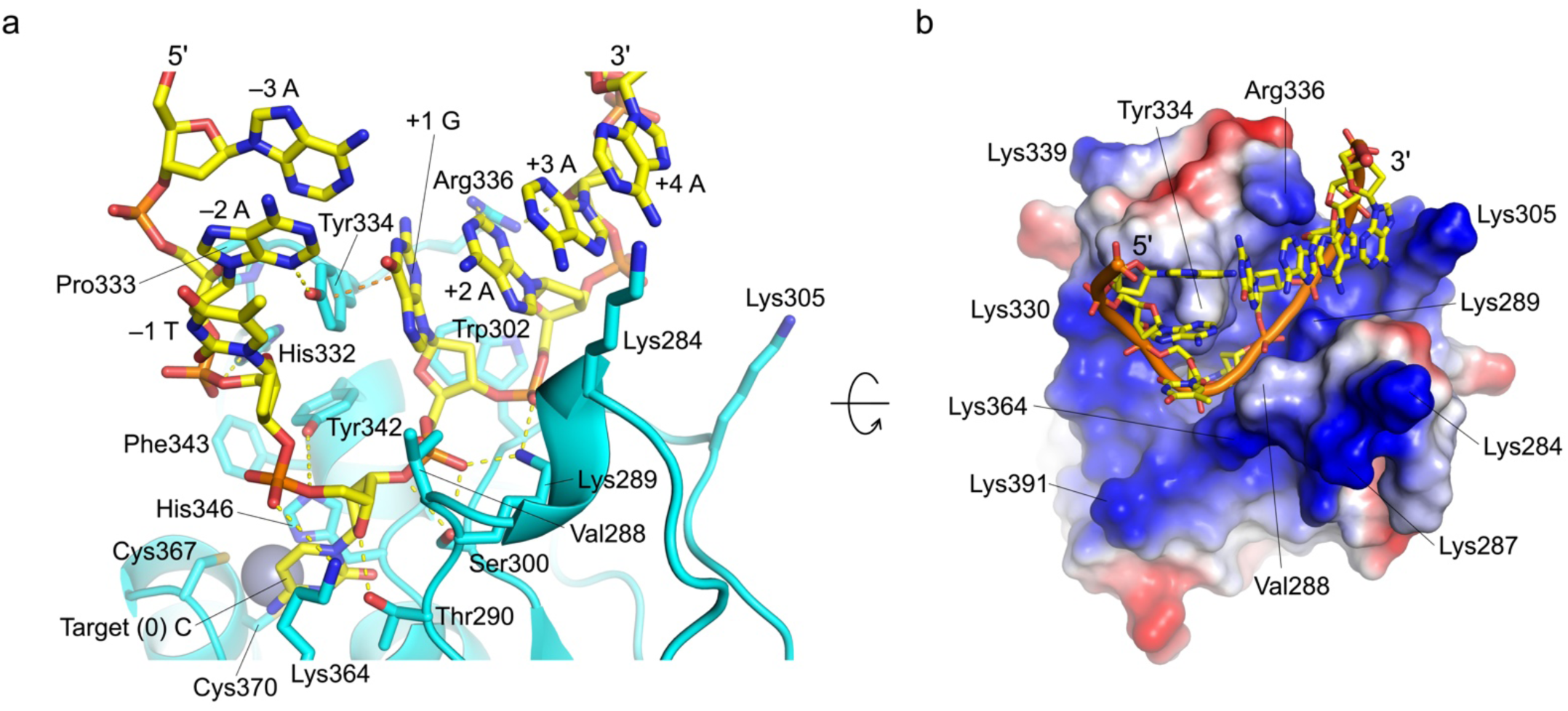
The mechanism of ssDNA substrate engagement by SsdA. **a**, Close-up view of the SsdA-ssDNA interaction, with hydrogen bonds and salt bridges indicated by yellow dashed lines. An orange dashed line indicates an aromatic stacking interaction between Tyr334 and +1 G. **b**, An orthogonal view looking down the narrow DNA-binding cleft, with the molecular surface of the protein colored according to electrostatic potential (blue: positive, red: negative).

In the deaminase active site, Thr290 plays a key role in poisoning the target (0) 2’-deoxycytidine nucleotide by stacking against the cytosine base and hydrogen-bonding to the deoxyribose moiety. The adjacent residues Lys289 and Ser300 interact with the phosphate backbone of +1G and +2A downstream of the target C. On the other side of the ssDNA-binding groove, Trp302 makes extensive van der Waals contact with the deoxyribose moieties of +1G and +2A, whereas Arg336 is hydrogen-bonded with the base and deoxyribose moieties of +2A and +3A, respectively, to stabilize the stacked nucleotides downstream of the target C.

A superposition of the antitoxin (SsdAI)-bound and the ssDNA-bound SsdA structures shows that SsdAI engages the tall loops on either side of the ssDNA-binding cleft and inserts Asp150 and Glu152 side chains into the ssDNA-binding cleft, which mimic the backbone phosphate groups of the nucleotides at +2 and +1 positions (**Fig. 3** and **Supplementary Fig. S7**). Furthermore, Tyr334 of SsdA fits in a hydrophobic pocket of SsdAI and makes a T-shaped ρε-stacking with an antitoxin residue Phe154, which takes the position of the +1G base of the ssDNA. Thus, as observed for DddI inhibition of DddA, SsdAI mimics both the shape and charge of the DNA substrate to achieve competitive inhibition with high affinity^3,7^.

**Figure 3.**
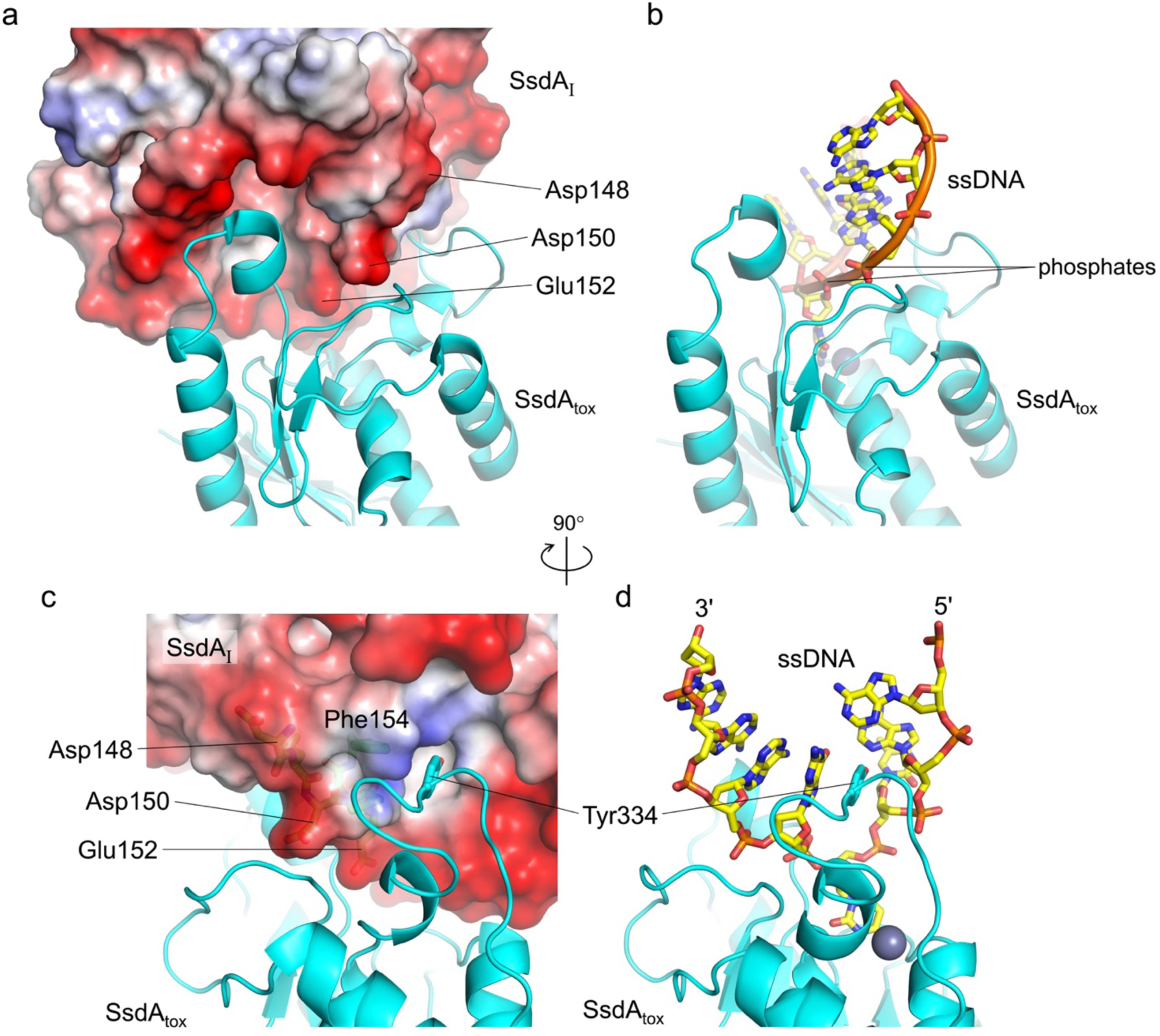
DNA mimicry by the anti-toxin, SsdAI. **a**, Crystal structure of the SsdAI-SsdAtox complex (PDB ID: 7jtu)^2^. The molecular surface is shown for SsdAI, colored according to electrostatic potential (blue: positive, red: negative). The side chains of acidic residues positioned similarly to DNA backbone phosphates are labeled. **b**, SsdAtox-DNA complex in the same orientation as **a**. **c**, Structure of the SsdAI-SsdAtox complex as in **a**, after a 90° rotation. Some of the SsdAI residues are shown in sticks and visible through a semi-transparent surface. **d**, SsdAtox-DNA complex in the same orientation as **c**.

### Mutation analyses reveal key aromatic contacts

To complement the crystallographic observations above and to identify functionally important residues, we examined the ssDNA deaminase activity of various SsdA mutant derivatives in a biochemical assay (**Fig. 4**). Consistent with the key role played by Tyr334 in ssDNA engagement, SsdA Y334A showed no detectable activity. In contrast, SsdA Y334F retained a near wildtype-level activity, highlighting the importance of an aromatic interaction. On the other hand, SsdA Y342A had no activity whereas SsdA Y342F was only weakly active. Similarly, a series of SsdA mutant derivatives H332A, H332Q, and H332T all showed no activity, whereas H332F showed only very weak, albeit detectable, activity. These results suggest that both the aromatic and polar interactions made by His332 and Tyr342 are important. P333A showed only a modest defect, indicating that Pro333 is less important than the neighboring residues His332 and Tyr334. A mutation of another residue Phe343 from the aromatic cluster, F343A, almost abrogated the deaminase activity. This suggests that Phe343, although it does not directly interact with ssDNA, plays a critical structural role.

**Figure 4.**
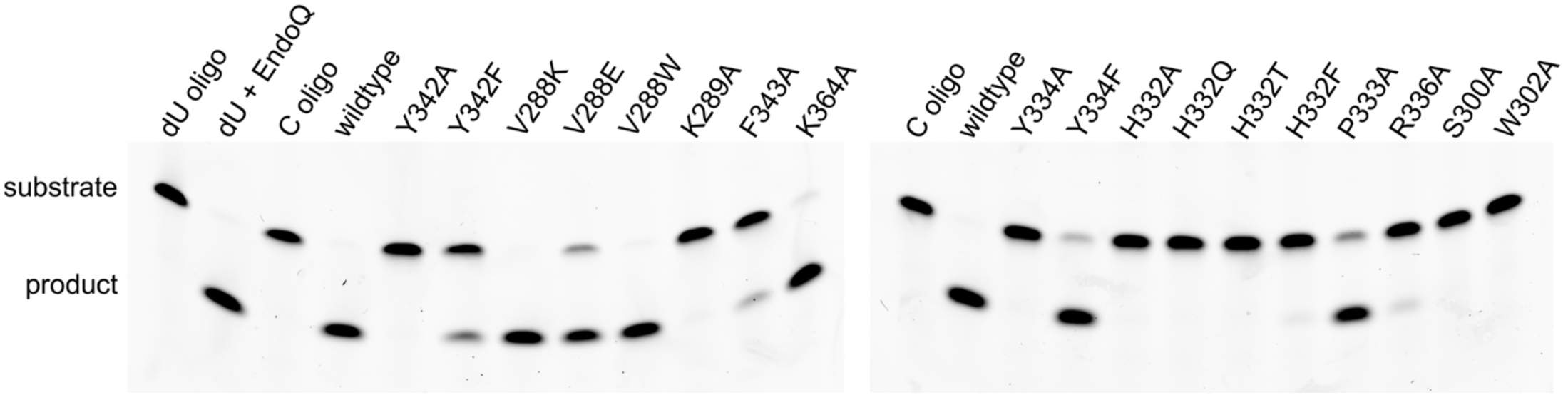
DNA cytosine deaminase activity of SsdA mutant derivatives. Deamination of ssDNA substrate (C oligo: FAM-5’-AAAAAAAAT<u>C</u>GAAAAAAA-3’, target cytosine underlined) by SsdA with various single amino acid substitutions for DNA-interacting residues around the active site (Fig. 2). Deaminated ssDNA oligonucleotide was cleaved by EndoQ^42^, yielding a band with faster mobility in gel electrophoresis. A chemically synthesized deamination product (dU oligo, far left) was confirmed to be fully cleaved by EndoQ (2nd lane).

SsdA K289A, S300A, W302A, and R336A showed no or little activity, underscoring the importance of these mutated residues interacting with nucleotides downstream of the target C. Notably, Gly301 takes a backbone conformation only allowed for glycine (β = ∼120°, ϕ ϑ = ∼160°) to enable the unique positioning of Ser300 and Trp302 for ssDNA interaction, and this ‘SGW motif’ is among the most highly conserved residues in BaDTF2 members (**Supplementary Figs. S1** and **S2**). For Val288 positioned near the mouth of the active site, substituting a positively charged (Lys) or a bulky aromatic (Trp) residue did not impair the SsdA activity, whereas an acidic (Glu) substitution at this position made SsdA slightly less active. Unexpectedly, SsdA K364A retained a near wildtype-level activity, suggesting that Lys364, despite its direct hydrogen bond with the 5’ phosphate of the target C, is functionally less critical.

### SsdA contains a β-amino acid

During the model building and refinement of the SsdA-ssDNA complex structure, it became apparent that the electron density for Asn294 could not be accounted for by the expected L-Asn and is more consistent with its β-linked isomer (**Supplementary Fig. S5**). To obtain stronger structural evidence, we crystallized SsdA without DNA and determined the structure at 1.33-Å resolution (**Table 1**; **Supplementary Fig. S6**). The high-resolution electron density for this structure unambiguously showed that residue 294 of SsdA is a β-amino acid in both molecules in the asymmetric unit (**Fig. 5a, b**). The side chain of residue 294 forms bidentate hydrogen bonds in an idealized geometry with Arg272 from a neighboring α-helix. This suggests that the residue is L-β-Asp or isoaspartate (isoAsp), which is known to result from a spontaneous deamidation of L-Asn via a succinimide intermediate^16^. IsoAsp294 is in a tight turn between short β-strands, where its side chain also accepts a hydrogen bond from the preceding residue Ser293 and its main chain amide nitrogen donates a hydrogen bond to the carbonyl oxygen of Lys296. Thus, isoAsp at this position plays an important structural role in stabilizing the protein fold. It is notable that isoAsp294 of SsdA is followed by Asp295, and this sequence context deviates from the more readily deamidated Asn-Gly and Asn-Ser motifs^17–19^, suggesting that the isomerization of SsdA Asn294 to isoAsp is conformation-specific.

**Figure 5.**
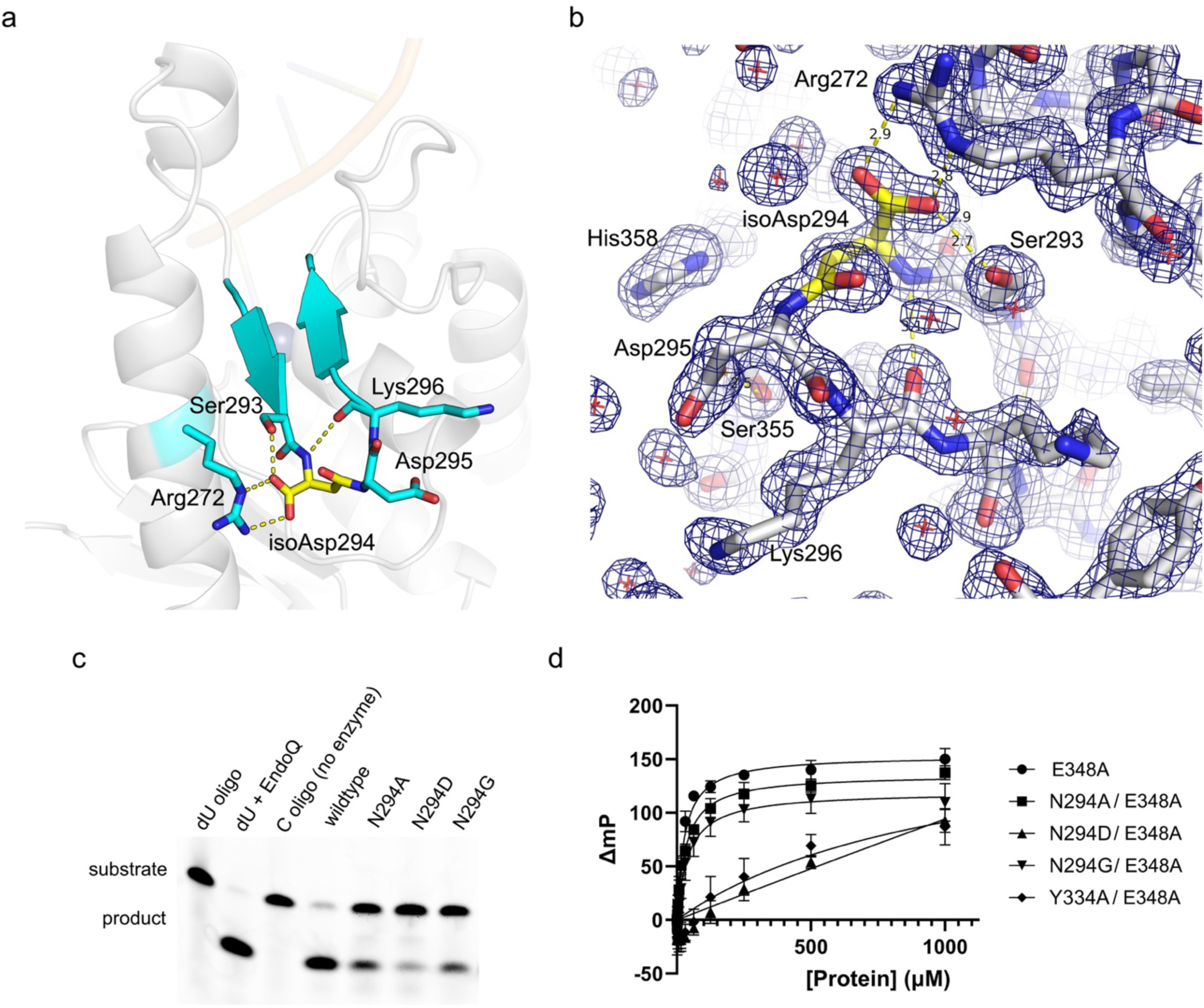
Isoaspartate (isoAsp) contributes to SsdA DNA-binding and deaminase activities. **a**, Crystal structure of the SsdA-ssDNA complex, with a β-turn containing isoAsp294 and the surrounding residues highlighted. Hydrogen bonds involving isoAsp294 are indicated by yellow dashed lines. **b**, The composite omit 2Fo-Fc electron density contoured at 1.5α for the higher-resolution SsdA structure without DNA. Hydrogen bonds involving isoAsp294 are indicated by yellow dashed lines, with distances in angstroms. c, DNA cytosine deaminase activity of SsdA mutant derivatives with residue 294 changed to Ala, Asp, or Gly. The enzyme concentration in this experiment was 16.5 nM. d, ssDNA-binding of SsdA mutant derivatives analyzed in a fluorescence polarization assay. All proteins carried the E348A mutation to inactivate the DNA deaminase activity. Y334A removes the key aromatic side chain of Tyr334 essential for the SsdA activity (Figs. 2 and 4).

To investigate the importance of isoAsp294 in the SsdA function, we tested the effect of N294A, N294D, and N294G mutations on SsdA activities. The alanine or glycine substitution should block the spontaneous post-translational isomerization to a β-amino acid. In addition, as Gly is achiral, Gly294 may better mimic the unique backbone conformation adopted by isoAsp294. The biochemical DNA deaminase assay showed that all 3 mutant derivatives are less active than wildtype SsdA, with N294A and N294G being more active among the mutant series, and N294D showing the most compromised activity (**Fig. 5c**). Although L-Asp could form the succinimide intermediate and go through the isomerization, L-Asp294 of SsdA was likely not converted into isoAsp. The effects of the three mutations were further evaluated for ssDNA-binding in a fluorescence polarization assay (**Fig. 5d**). The crystallized protein SsdA E348A showed a KD of 28.0 μM for a 17-nt ssDNA containing a TCG motif. A double mutant Y334A/ E348A, which has the key DNA-binding residue Tyr334 mutated, exhibited a much weaker binding with KD > 378 μM (95% CI). SsdA N294D/E348A showed even poorer binding (KD not determined due to the lack of saturation in binding). In contrast, SsdA N294A/E348A and N294G/E348A exhibited comparable affinities (KD of 33.6 and 42.3 μM, respectively) to SsdA E348A, consistent with what we observed in the DNA deamination assay above that substitution of Ala or Gly had milder effects than Asp at residue 294. These results suggest that, while isoAsp294 is not essential for substrate engagement, it contributes to optimal ssDNA substrate-binding and deamination activities of SsdA.

### SsdA mutant derivatives with activity in human cells

The sequence context-independent ssDNA deaminase activity of SsdA could be useful in base editing or other applications in human cells^20^. Thus, we tested the cytosine deaminase activity of SsdA in 293T cells using the real-time, live cell-based ARSENEL assay against four different dinucleotide sequence contexts^21,22^. Wildtype SsdA fused to Cas9n (D10A) in the BE4max platform^23^ showed activity with all four dinucleotide target motifs (AC/CC/GC/TC), although the editing efficiency on the GC target is significantly lower (**Fig. 6a**). Curiously, a series of biochemically less active mutant derivatives of SsdA: N294A, N294D, and N294G, all showed significantly higher editing efficiencies than wildtype SsdA, especially for the GC target. Immunoblot analysis confirmed that the SsdA-based editors are expressed, albeit at lower levels compared to the APOBEC3A (A3A)-based editor used as a control, likely due to cytotoxicity (**Fig. 6b**). These results demonstrate that SsdA is active in human cells and suggest that mutant derivatives may be promising candidates for use in base editing and other applications. Notably, a recent study used SsdA in base editing in mammalian cells including mouse embryos^24^, consistent with our observations.

**Figure 6.**
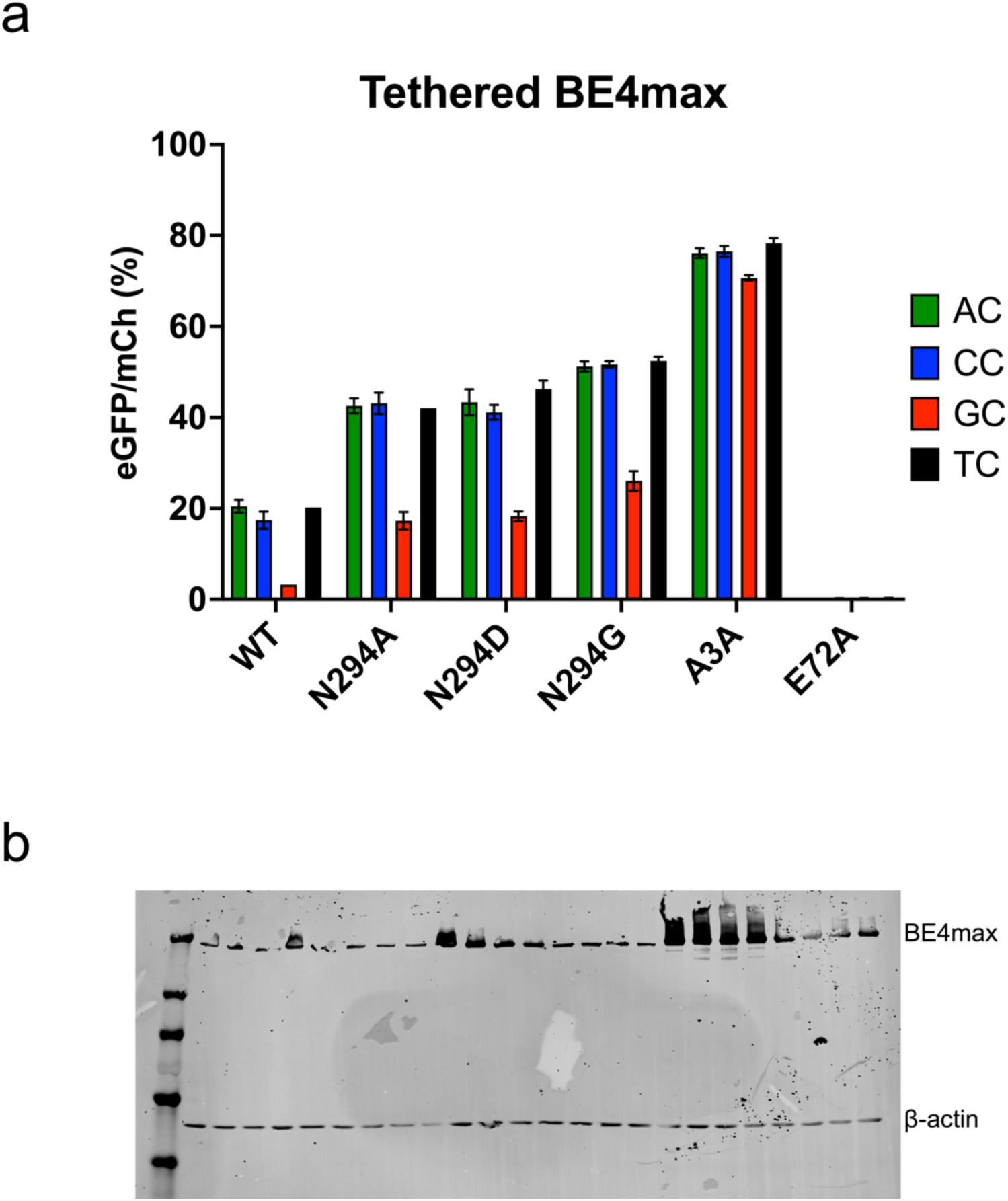
SsdA-mediated base editing in human cells. **a,** Base editing efficiencies of A3A- and A3A-E72A-BE4max in comparison to wildtype (WT) SsdA- Cas9n or the indicated mutant variants quantified using flow cytometry. Editing efficiencies were assessed on four distinct ARSENAL reporter constructs following co-transfection of all ARSENEL assay components into 293T cells. Column bars represent the mean ± SD from biologically independent triplicate experiments. **b,** Immunoblot analysis of base editor expression using an anti- Cas9 antibody (lane order matches constructs in bar graph in panel a). β-actin was used as a loading control.

## Discussion

SsdA shapes ssDNA into a unique V-shape to engage the target cytosine in the active site. The sharply bent ssDNA substrate, with the target deoxycytidine nucleotide flipped out for an engagement in the zinc-bound active site, is a shared feature with the APOBEC-family ssDNA deaminases.

However, a superposition of the SsdA-ssDNA complex and APOBEC3A-ssDNA complex^25^ based on the conserved core structural elements of the deaminase fold shows distinct trajectories of the ssDNA backbones, with the DNA strands approaching the active site in orthogonal orientations. In the APOBEC3A/B-DNA complex, both the target (0) deoxycytidine and the 5’ neighboring (–1) deoxythymidine nucleotides are rotated out at the bottom of a ‘U’-shaped ssDNA and engaged by the enzyme through base-specific contacts, underlying the strong 5’-TC preference^25^. In contrast, in the SsdA-bound ssDNA, only the target deoxycytidine is rotated out toward the protein at the apex of the V-shape and makes sequence-specific contact, which explains the lack of a strong –1 base preference by SsdA (**Fig. 7a** and **Supplementary Fig. S4**).

**Figure 7.**
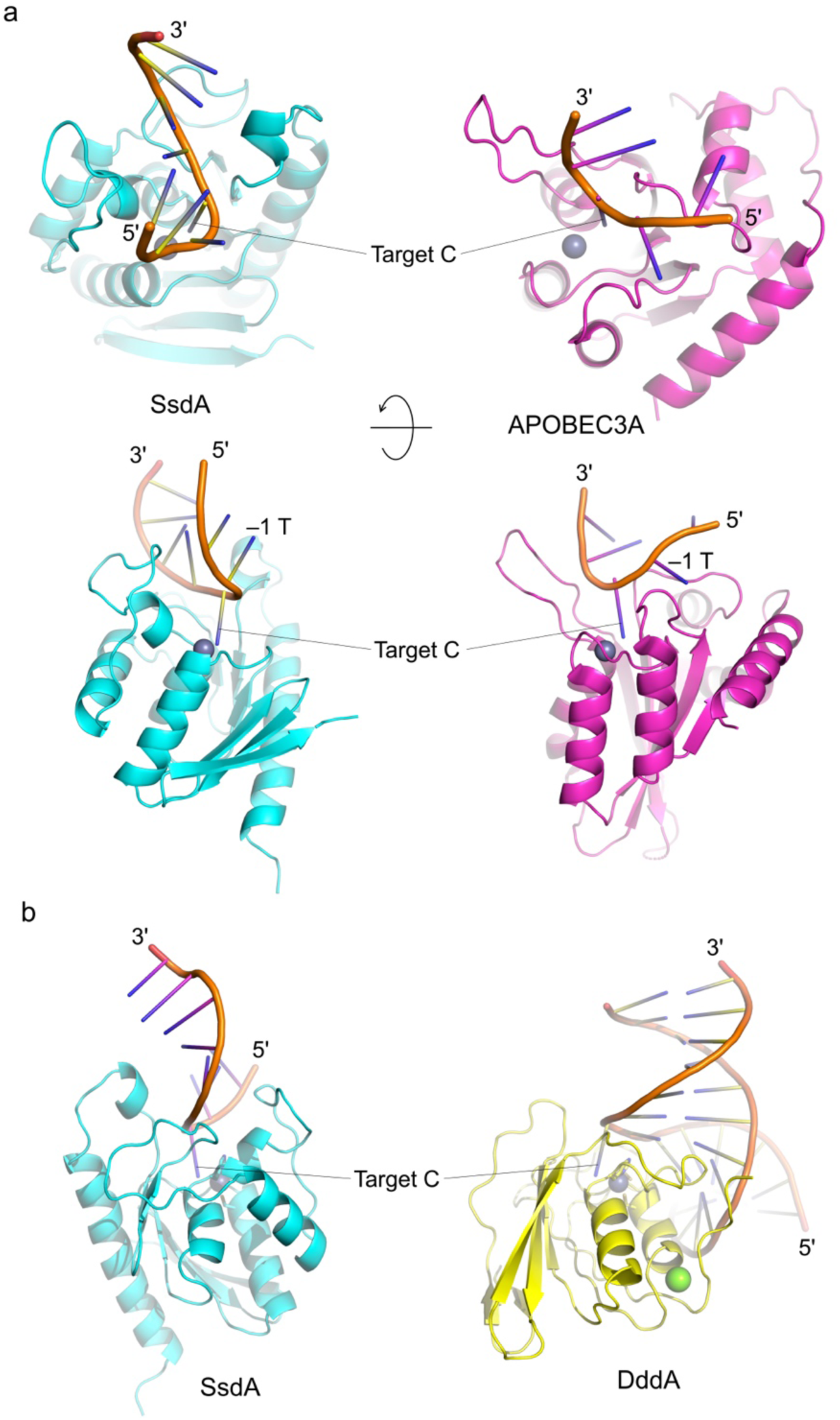
Modes of target engagement by SsdA and related DNA deaminase enzymes. **a,** Side-by-side comparisons between the SsdA-ssDNA complex structure (cyan, left) and the human APOBEC3A-ssDNA complex structure (magenta, right) (PDB ID: 5SWW)^25^. In each (upper and lower) pair, the two enzymes are shown in the same orientation based on a superposition of the conserved active site residues. The gray spheres represent a Zn ion in the active site. **b,** A side-by-side comparison between the SsdA-ssDNA complex structure (cyan, left) and the DddA-dsDNA complex structure (yellow, right) (8E5E)^7^. The two enzymes are shown in the same orientation based on a superposition of the conserved active site residues. The gray spheres represent a Zn ion in the active site. The green sphere bound to DddA is a magnesium ion.

A structural comparison between the SsdA-ssDNA complex and the DddA-dsDNA complex^7^ highlights a remarkable diversity of DNA-binding mode among BaDTFs (**Fig. 7b**). The core of the deaminase fold comprising two α-helices packed against a β-sheet is sandwiched between additional α-helices in SsdA, which support the extended loops on either side of the deep ssDNA-binding cleft. In contrast, the core of DddA is flanked on either side by loops making smaller protrusions, which together form a broader DNA-binding surface for dsDNA substrates. Interestingly, BaDTFs don’t show exquisite selectivity for dsDNA or ssDNA^10^, and SsdA shows significant activity on dsDNA substrates (**Supplementary Fig. S4**). A superposition between the two protein molecules in the asymmetric unit of the high-resolution DNA-free SsdA crystal structure shows the largest deviation in a loop centered on Lys284, suggesting the flexibility of this region (**Supplementary Fig. S8**). This loop faces the base-pairing edges of the linearly stacked nucleobases in the SsdA-bound ssDNA, and thus, the flexibility might allow SsdA to accommodate dsDNA or partially melted dsDNA as a substrate.

Deamidation of Asn into isoAsp in proteins is generally considered spontaneous damages that negatively affect their stability and activity^17,26^. IsoAsp is recognized by protein L-isoaspartyl methyltransferase (PIMT), an enzyme conserved between prokaryotes and eukaryotes that methylates isomerized or racemized aspartyl residues to facilitate their conversion back to canonical L-Asp^27^. However, isoAsp in special cases has been reported to confer advantageous properties in bacterial, eukaryotic, and viral proteins^28–30^. For instance, an essential enzyme MurA from *Enterobacter cloacae* contains an isoAsp residue in a β-hairpin turn, which contributes to protein stability^31^. IsoAsp294 of SsdA is similarly in a hairpin turn that supports the positioning of DNA-interacting loops and thus contributes to the enzymatic activity. It is possible that isoAsp also serves to confer resistance against proteolytic degradation in target cells and thereby enhances the effectiveness of SsdA as an interbacterial toxin. Conversely, this post-translational modification could conceivably make SsdA susceptible to inactivation by the PIMT activity in receiving bacteria.

Regardless, the efficient base-editing activity of SsdA N294A/D/G in human cells and the structural information presented here provide a basis for the rational design of engineered DNA deaminases for novel applications.

## Materials and Methods

### Protein expression and purification

SsdA (residues 259-409) catalytically inactive mutant E348A, and double mutants N294A/E348A, N294G/E348A N294D/E348A, and Y334A/E348A were expressed in the *E. coli* strain BL21(DE3) using pET-24a expression vector with a C-terminal His6-tag. Transformed bacteria were grown in 4 liters of LB medium to an OD600 of 0.8, at which point the culture was supplemented with 100 μM ZnCl2 and induced by the addition of Isopropyl β-D-1-thiogalactopyranoside at a final concentration of 0.5 mM. After 20 hrs of shaking and incubation at 18°C, the bacterial cells were collected by centrifugation at 4,000xg for 30 min. The cell pellets were resuspended in 20 mM Tris-HCl, pH 7.4, 0.5 M NaCl, 5 mM β-mercaptoethanol and 5 mM imidazole, lysed by the addition of lysozyme (0.4 mg/ml) and sonication. The lysate was centrifuged at 64,000xg for 1 hr at 4°C to separate the supernatant from cell debris. The supernatant was collected and filtered through a 0.2 μm asymmetric polyethersulfone membrane and applied to a 5 mL Ni-NTA Superflow cartridge (QIAGEN). The His-tagged SsdA protein was eluted with a linear concentration gradient of imidazole and further purified by size-exclusion chromatography (SEC) on a HiLoad 26/600 Superdex 75 column operating with 20 mM Tris-HCl, pH 7.4, 0.2 M NaCl, 0.5 mM tris(2-carboxyethyl)phosphine (TCEP). Purified proteins were concentrated by ultrafiltration using Amicon centrifugal filters (3 kDa MWCO). The protein concentrations were measured on a Nanodrop 8000 spectrophotometer based on UV absorbance at 280 nm and the extinction coefficients calculated from the amino acid sequences.

### X-ray crystallography

SsdA E348A was mixed with a 1.5-fold molar excess of 15 nt ssDNA (5’-AAAAAATCGAAAAAA-3’) in 10 mM Tris-HCl pH 7.4, 100 mM NaCl, and 0.5 mM TCEP at the final protein concentration of 27.5 mg mL^-1^. The SsdA-DNA complex was crystallized by the hanging drop vapor diffusion method using a reservoir solution containing 0.15 M NH4Cl and 22.5% PEG3350, pH 6.3, and the 1:1 volume ratio of mixing the protein-DNA complex and reservoir solutions in forming the drop. The DNA-free SsdA E348A crystals were obtained similarly, by mixing the protein at 55 mg mL^-1^ with a reservoir solution containing 1.8 M sodium phosphate monobasic monohydrate and potassium phosphate dibasic, pH 7.8. The crystals were cryo-protected by brief soaking in the respective reservoir solution supplemented with 20% ethylene glycol and flash-cooled by plunging in liquid nitrogen. X-ray diffraction data were collected at the NE-CAT beamline 24-ID-E of the Advanced Photon Source (Lemont, IL) for the SsdA-ssDNA crystal, and at the AMX beamline of NSLS-II (Upton, NY) for the DNA-free SsdA crystal. All datasets were processed with XDS^32^ for integration, followed by three other programs from the CCP4 Suite^33^: POINTLESS^34^, AIMLESS^35^ and TRUNCATE^36^ for reduction, scaling, and structure factor calculation, respectively. Anisotropic diffraction analysis and truncation were done with STARANISO (https://staraniso.globalphasing.org/). The structures were determined by molecular replacement with PHASER^37^ using the previously reported crystal structure of SddAtox bound to SddAI (PDBID 7JTU)^2^ as the search model. Iterative model building and refinement were conducted using COOT^38^ and PHENIX^39^. Modeling an L-Asn at residue 294 according to the genetic sequence renders the N294 side chain out of the density and results in a distorted geometry of the nearby backbone and strong positive and negative Fo-Fc densities. However, modeling the 294 position as isoAsp generates good geometry and clears the Fo-Fc map around the 294 position. A summary of crystallographic data statistics is shown in **Table 1**. Figures were generated using PyMOL (https://pymol.org/).

### Cell-free protein expression and purification

Due to the toxicity of SsdA in bacterial cells, wildtype SsdA and various mutant derivatives with single amino acid substitutions used in the biochemical assay were expressed using NEBExpress® Cell-free *E. coli* Protein Synthesis System (New England Biolabs). Linear dsDNA templates used for expression were chemically synthesized as gBlock DNA fragments by Integrated DNA Technologies. The sequences of the gene fragments were confirmed by Sanger sequencing (ACGT). The cell-free expression mixture was incubated at 37°C for 2.5 hrs and flash-frozen in liquid nitrogen to stop the reaction. His-tagged wildtype or mutant SsdA was affinity-purified with Dynabeads™ His-Tag Isolation and Pulldown kit (ThermoFisher Scientific). The purified proteins were confirmed by SDS-PAGE and their concentrations were quantitated based on band intensities in SDS-PAGE using Image J^40^ against previously quantitated SsdA E348A as a standard.

### Biochemical DNA deaminase assay

The ssDNA oligonucleotide used in deaminase assay is a 5’-fluorescein-labeled 18-nt DNA (FAM- 5’-AAAAAAAATCGAAAAAAA-3’) and its variants with different –1 nucleobases (5’-AC, 5’-CC, 5’- GC). The reaction mixture contained 200 nM ssDNA substrate, wildtype or mutant SsdA at 57 nM (or otherwise stated in figure legends), 40 mM Tris–HCl, pH 7.4, 50 mM KCl, and 1.0 mM dithiothreitol. After incubating at 37°C for 50 min, pfuEndoQ was added to the final concentration of 1.0 μM, and the samples were further incubated at 60 °C for 30 min to cleave deaminated products. The reactions were stopped by the addition of formamide to 65% and heating to 95 °C for 10 min. The products were separated by 15% denaturing PAGE and scanned on a Typhoon FLA 7000 imager (GE Healthcare). For each experiment, a control substrate representing the deaminated product (dU oligo) was treated with pfuEndoQ in parallel to confirm complete cleavage.

### Fluorescence polarization assay

The ssDNA-binding affinities of SsdA E348A, N294A/E348A, N294D/348A, N294G/348A, and Y334A/E348A were determined by fluorescence polarization/anisotropy measurements using a 3’-fluorescein-labeled 17-nt ssDNA probe (5’-GCGAAGTTCGGTTAACG-3’-FAM). A two-fold serial dilution series of protein concentrations ranging from 1.95 μM to 1.0 mM were mixed with the ssDNA probe at a final concentration of 18 nM in 20 mM Tris–HCl pH 7.4, 100 mM NaCl, 5 mM ϕ3- mercaptoethanol and the final volume 100 μL in a black flat-bottom half-area 96-well plate.

Changes in fluorescence polarization induced by protein binding were measured on a Tecan Spark 10M microplate reader. The data were fit to the one-site specific binding model in GraphPad Prism 9 to determine the KD values and 95% confidence intervals.

### ARSENEL assay

Subconfluent 293T cells in 12-well plates were transfected in parallel with 300 ng of each ARSENEL reporter, 100 ng of gRNA, and 300 ng of each base editor (20 min at room temperature with 1.8 μL of TransIT-LT1 (Mirus) and 100 μL of Opti-MEM (ThermoFisher Scientific). After 24 hrs of incubation, the transfected cells were PBS-washed and trypsinized for flow cytometry analysis using a BD LSRFortessa flowcytometer. A minimum of 100,000 events were collected for each condition, and the ratio of eGFP and mCherry double-positive cell number to the total number of mCherry-positive cells is calculated to indicate the editing efficiency.

### Immunoblotting

The remaining cells from flow cytometry experiments were collected for immunoblotting. The cells were PBS-washed and then lysed in RIPA buffer (ThermoFisher Scientific). A total of 24 samples were run on the same Criterion 4–20% gel (Bio-Rad), then transferred to a PVDF membrane.

Primary antibodies were β-Actin (13E5) Rabbit mAb (#4970, Cell Signaling Technology, 1:2,500), and SpCas9 Mouse mAb (#Ab191468, Abcam, 1:5000). Secondary antibodies used were goat anti-rabbit IRdye800 (LI-COR, #925-32211, 1:10,000) and goat anti-mouse IRdye680 (LI-COR, #926-69020, 1:10,000).

## Acknowledgments

ray diffraction data were collected at the Northeastern Collaborative Access Team beamlines of the Advanced Photon Source and the Center for Bio-Molecular Structure (CBMS) AMX beamline of NSLS-II, which are funded by the US National Institutes of Health grants NIGMS P30 GM124165 and P30 GM133893, respectively. This publication resulted from data collected during beamtimes obtained through NE-CAT BAG proposal #311950. This work was supported by NIH grants (NIGMS R35-GM118047 to H.A. and NCI P01-CA234228 to R.S.H. and H.A.) and a Recruitment of Established Investigators Award from the Cancer Prevention and Research Institute of Texas (CPRIT RR220053) to R.S.H. R.S.H. is the Ewing Halsell President’s Council Distinguished Chair at the University of Texas San Antonio and an Investigator of the Howard Hughes Medical Institute.

## Data Availability

Atomic coordinates and structure factors for the SsdA-DNA complex and SsdA without DNA have been deposited in the Protein Data Bank (PDB) under the accession codes 9C64 and 9C63, respectively. All other data are available from the authors upon request.

## Competing Interests

The authors declare no competing interests.

---

**Supplementary Figure S1.**
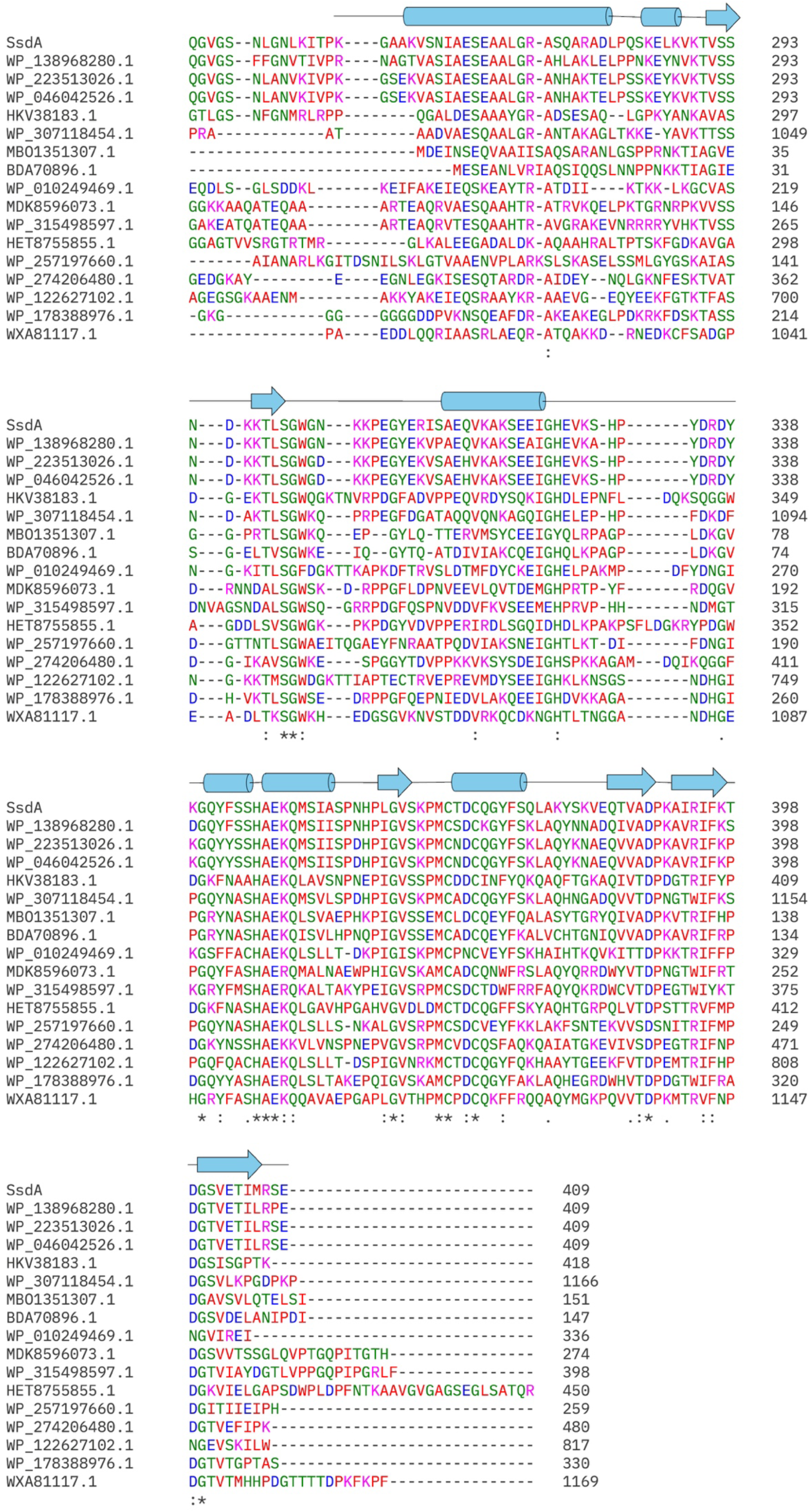
Amino acid sequence alignment of SsdA orthologs (BaDTF2) The sequence of the C-terminal deaminase domain of SsdA (SsdAtox) is at the top, with its secondary structure elements shown schematically above. The other sequences are identified by GenBank accession codes. The multiple sequence alignment was done using Clustal Omega^43^.

**Supplementary Figure S2.**
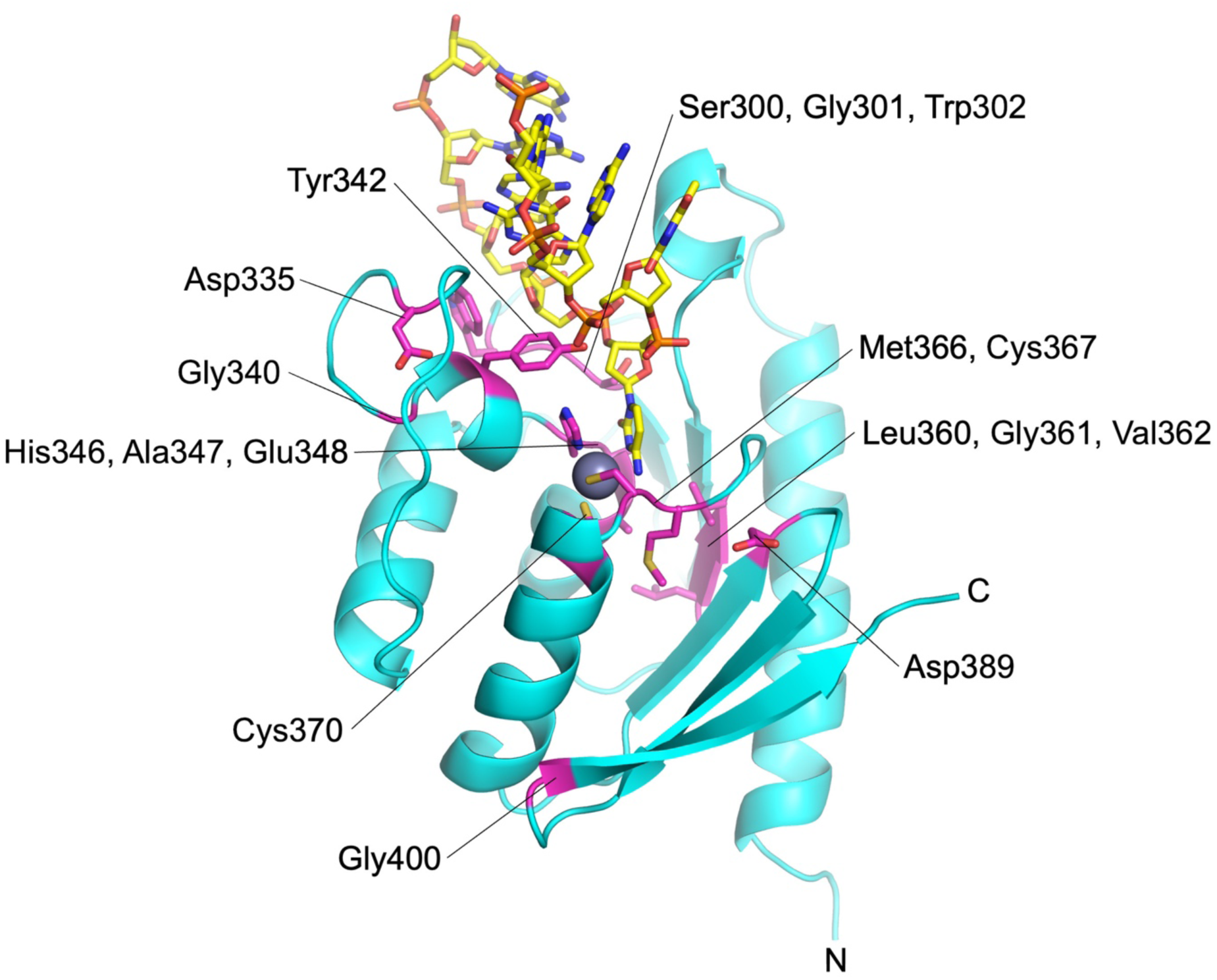
Highly conserved residues among the SsdA orthologs (BaDTF2) The positions of the residues showing high conservation in Supplementary Fig. S1 are highlighted in magenta, with their side chains shown in sticks. These residues mostly cluster around the ssDNA-binding cleft, suggesting a conservation of the general mode of ssDNA engagement.

**Supplementary Figure S3.**
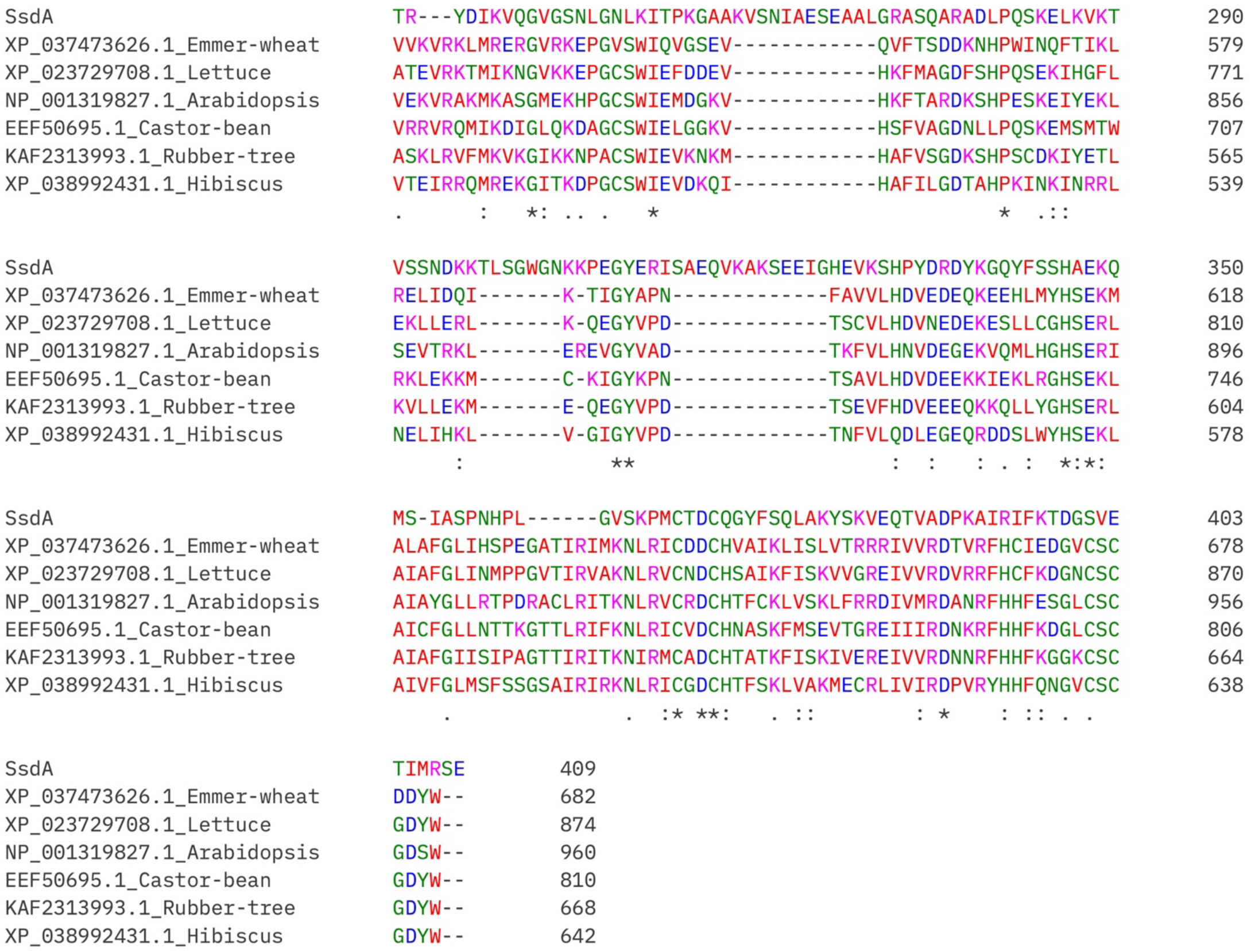
Amino acid sequence alignment between SsdA and DYW-family RNA deaminases. The sequence of the C-terminal deaminase domain of SsdA (SsdAtox) is at the top. The other sequences are identified by GenBank accession codes followed by a common name of the organism from which the sequence is derived. Despite high conservation of the catalytic residues, SsdA lacks the C-terminal DYW motif present in these plant RNA deaminases. The multiple sequence alignment was done using Clustal Omega^43^.

**Supplementary Figure S4.**
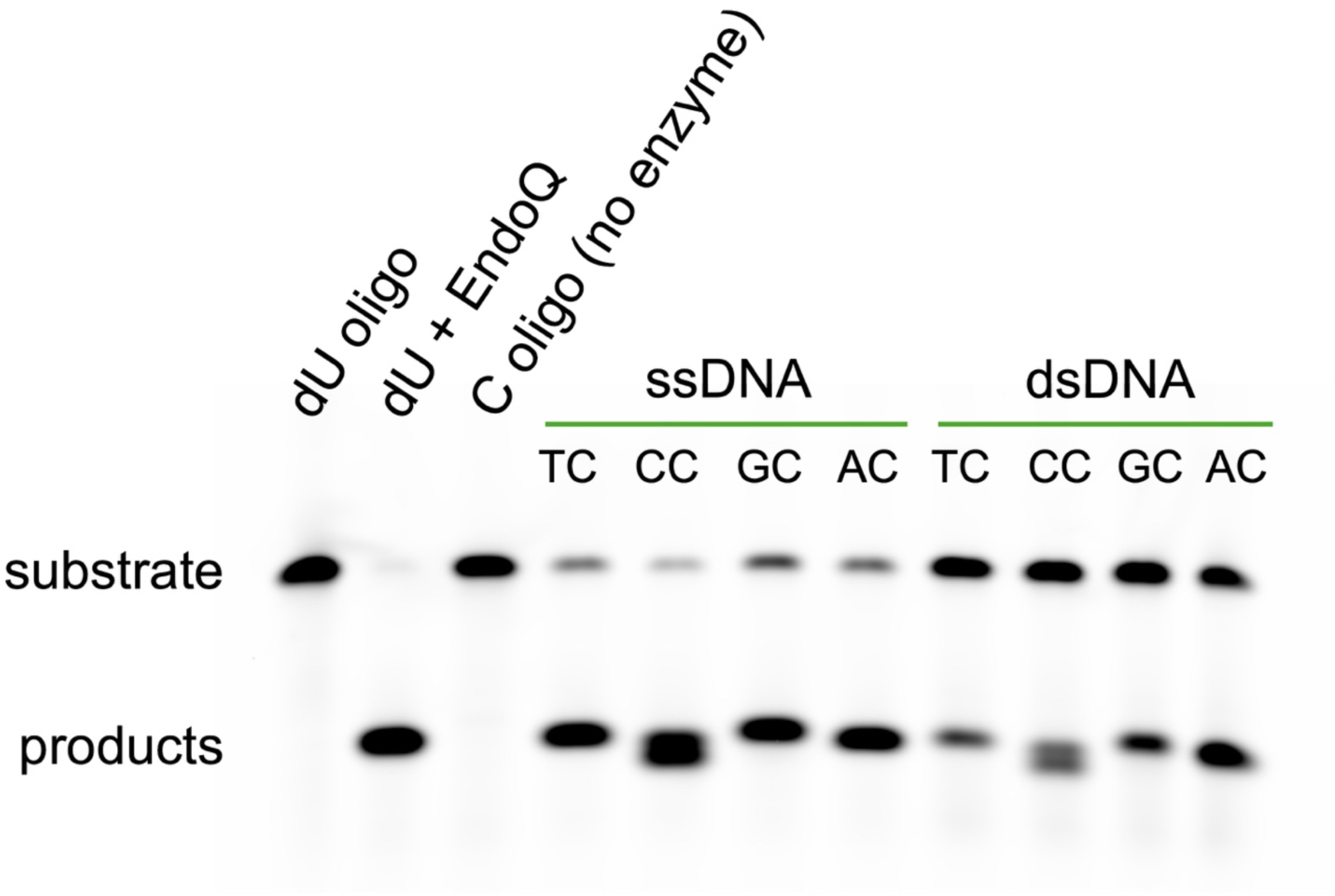
DNA substrate preferences of SsdA. Deamination by wildtype SsdA of the underlined cytosine base in a target oligo (FAM-5’- AAAAAAAAX<u>C</u>GAAAAAAA-3’, in which X is T, C, G, or A), either as ssDNA or annealed to the complementary strand as dsDNA, examined in an EndoQ-mediated oligo cleavage assay^42^. The SsdA concentration in this experiment was 8.2 nM. Note that the lower band (smaller product) for the CC substrate results from deamination at –1 C, which is in an ACC context.

**Supplementary Figure S5.**
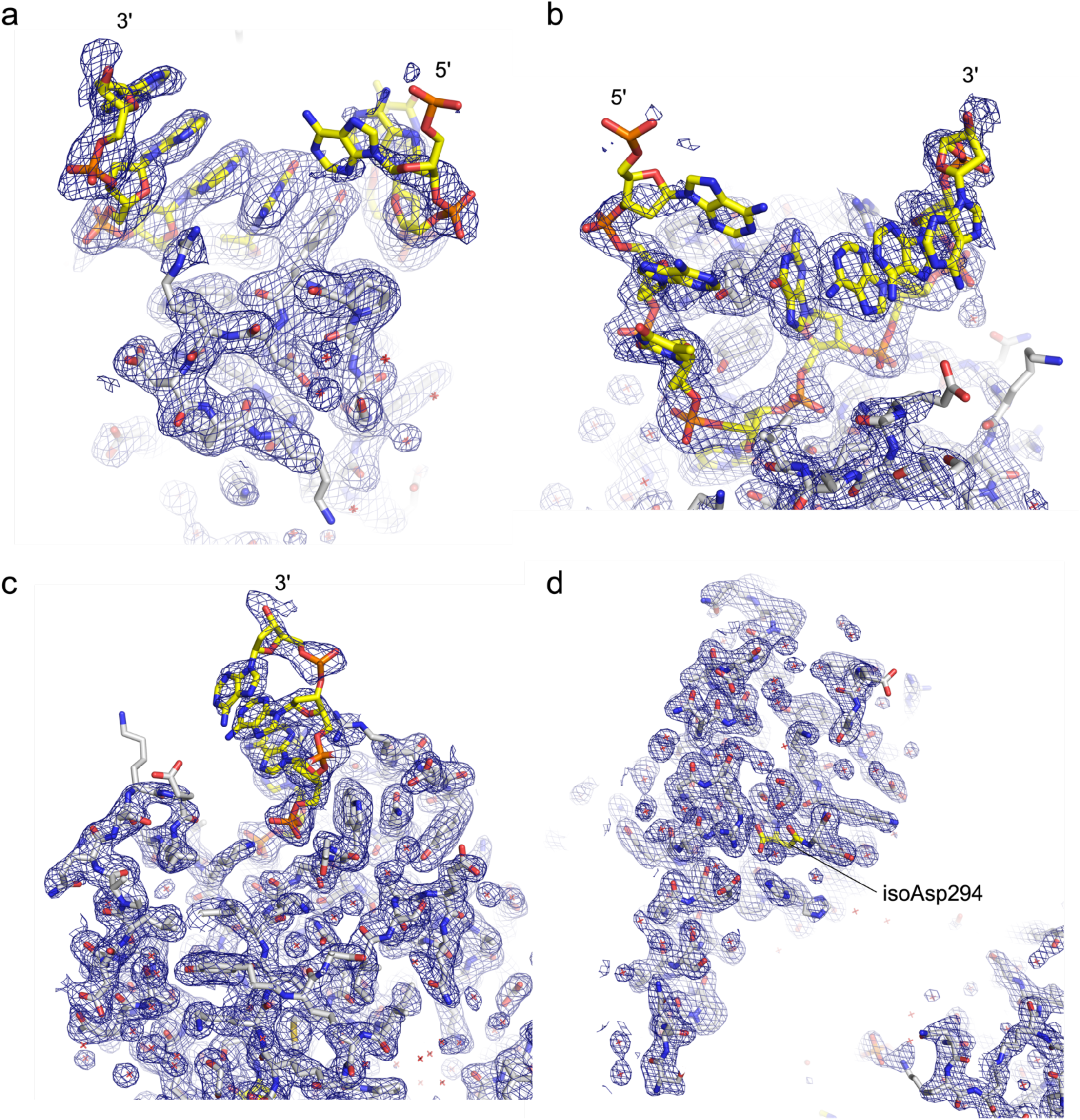
Electron density map for the SsdA-ssDNA complex structure. The composite omit 2Fo-Fc electron density contoured at 1.0σ is shown for various regions of the structure. The N-terminal long α-helix interacting with isoAsp294 (in yellow) is shown in **d**.

**Supplementary Figure S6.**
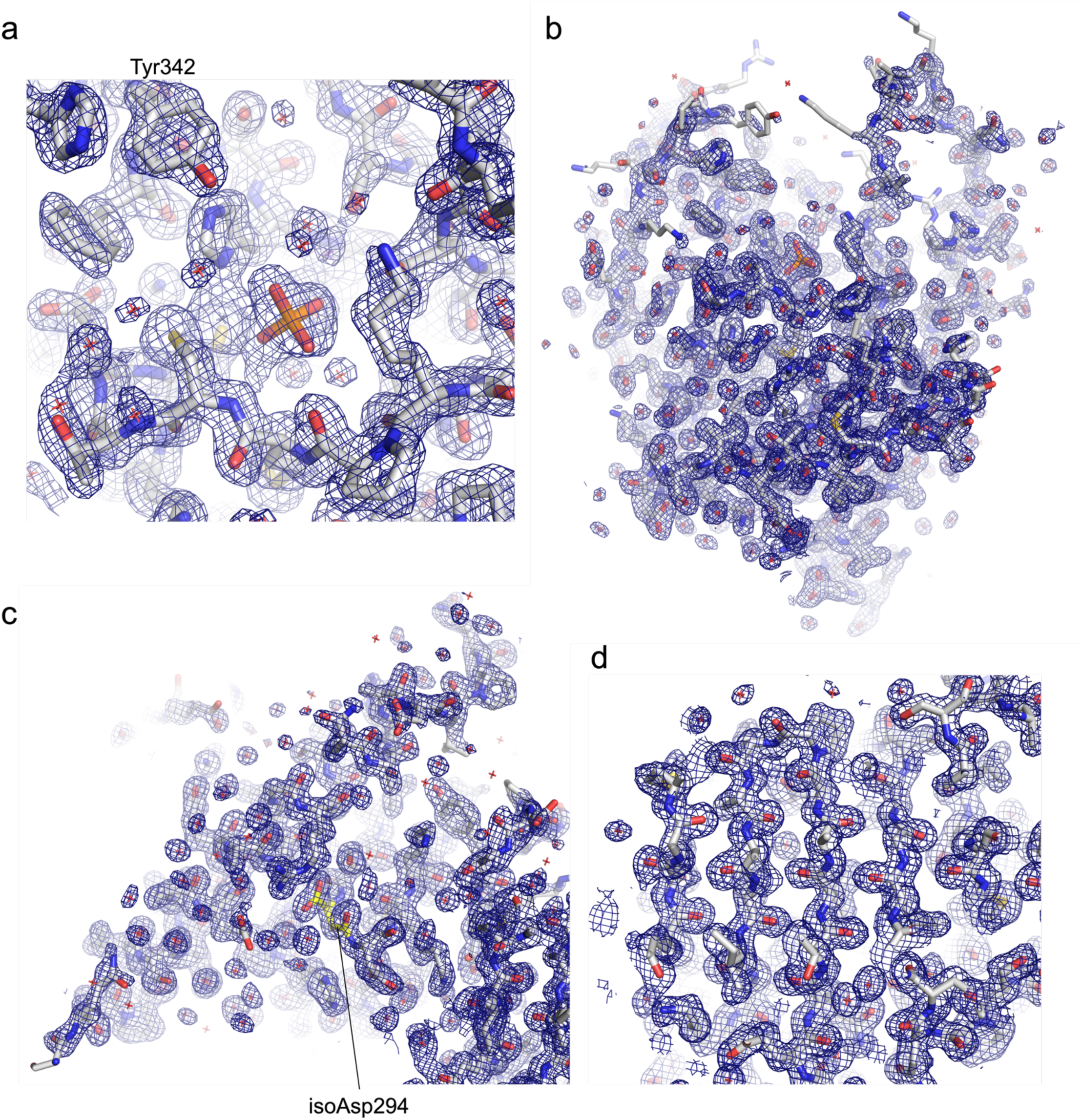
Electron density map for the DNA-free SsdA structure. The composite omit 2Fo-Fc electron density contoured at 1.0σ is shown for various regions of the structure. The active site (**a**) is occupied by a phosphate ion present in the crystallization condition.

**Supplementary Figure S7.**
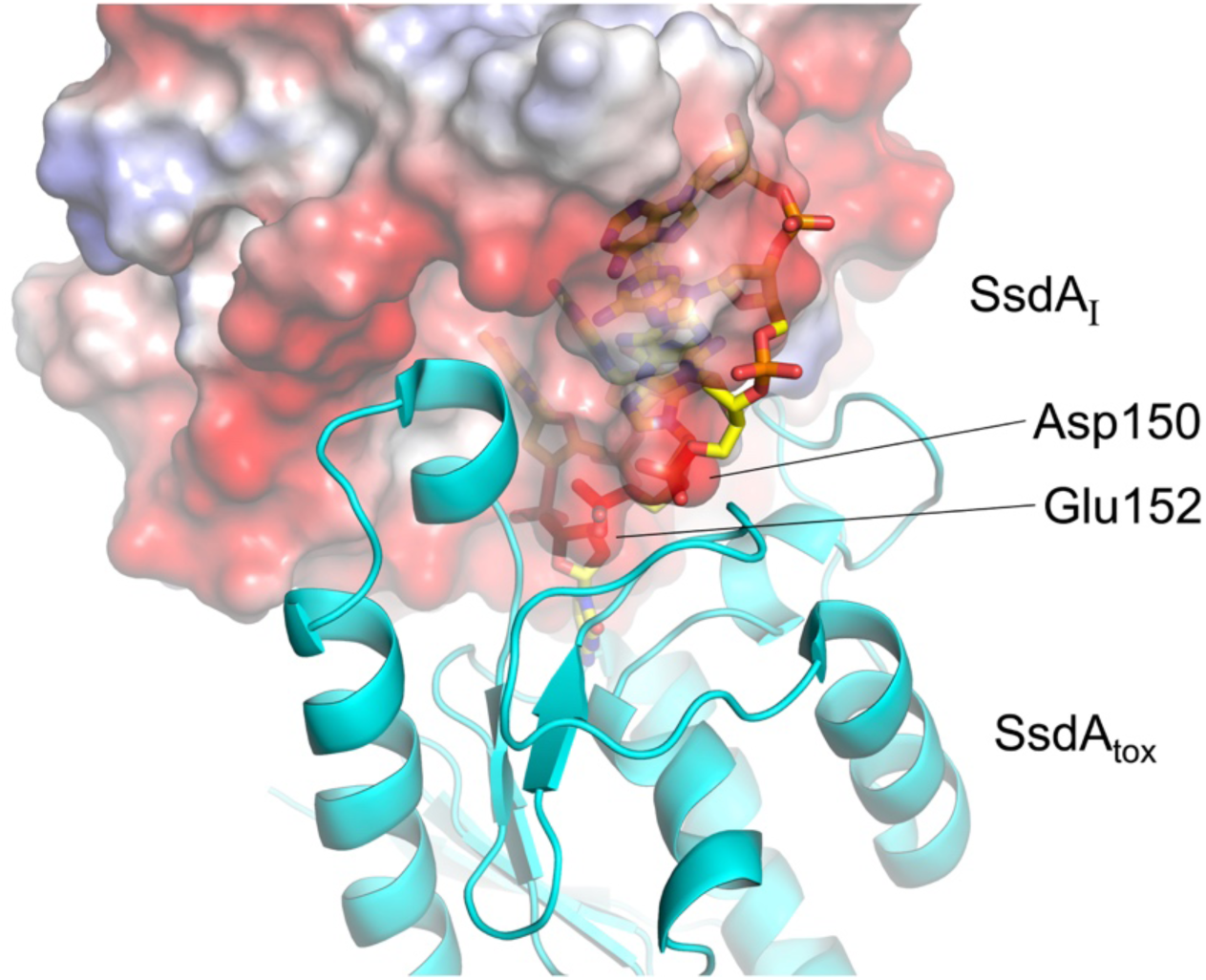
Mimicry of DNA backbone phosphates by SsdAI Asp150 and Glu152. An overlay of Fig. 3 a and b, highlighting the mimicry of DNA backbone phosphates by the negatively charged protein side chains.

**Supplementary Figure S8.**
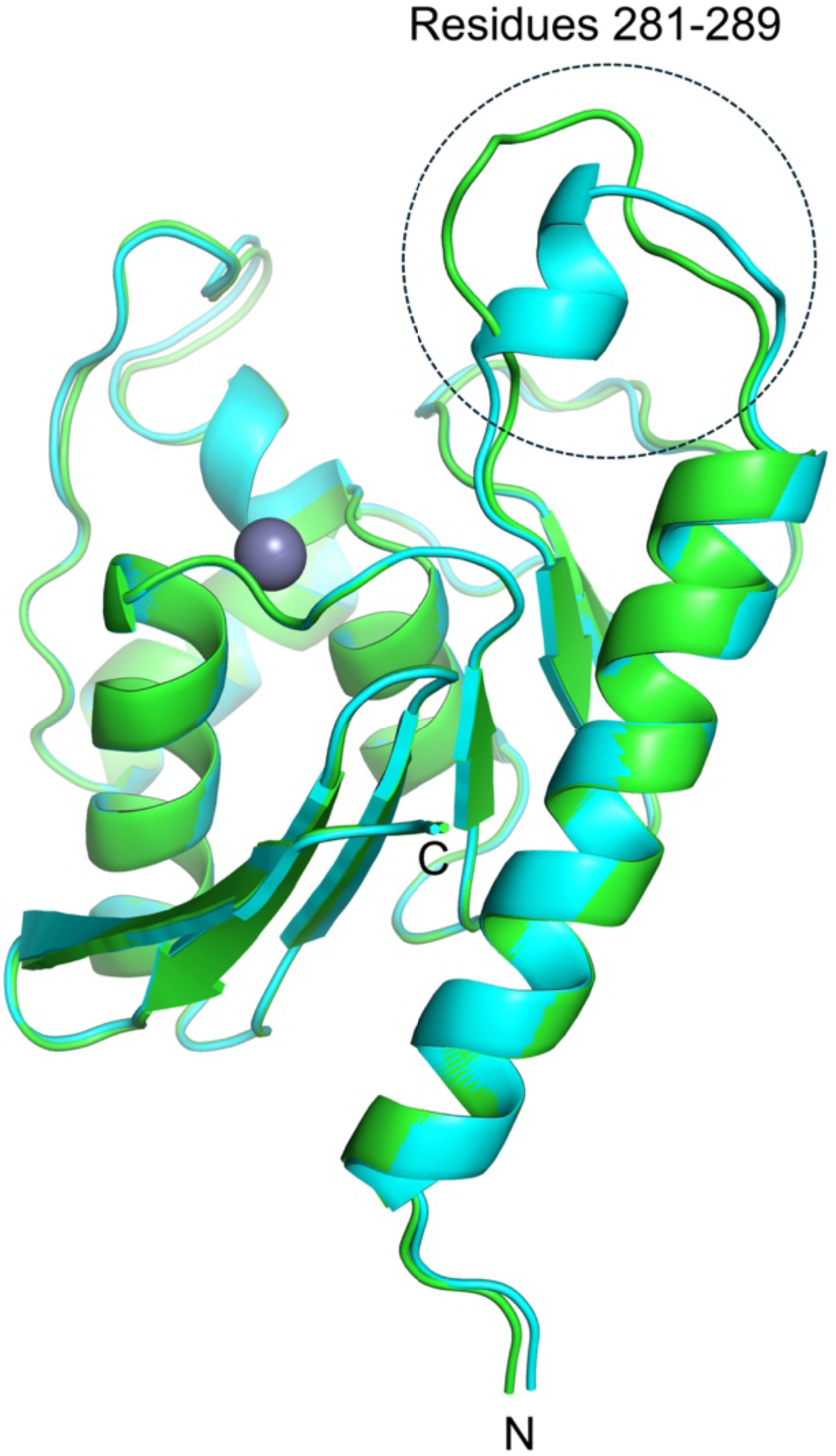
Structural plasticity in SsdA. A superposition between the two SsdA molecules in the asymmetric unit of the high-resolution, DNA-free SsdA crystal structure. The region that shows the greatest deviation is highlighted by a dashed circle.

## Notes

### Competing Interest Statement

The authors have declared no competing interest.

